# Improving AlphaFold 3 structural modeling by incorporating explicit crosslinks

**DOI:** 10.1101/2024.12.03.626671

**Authors:** Konstantin Gilep, Agnieszka Obarska-Kosinska, Jan Kosinski

## Abstract

AlphaFold 3 has significantly advanced the modeling of macromolecular structures, including proteins, DNA, RNA, and their interactions with small molecules or post-translational modifications. However, challenges remain when modeling specific structural conformations or complexes with limited evolutionary data, such as protein-antibody complexes. Previous studies with AlphaFold2 demonstrated that adding distance restraints from crosslinking mass spectrometry (XL-MS) can improve predictions for such cases. In this study, we investigate whether XL-MS restraints can be incorporated into AlphaFold 3 by explicitly modeling crosslinks as covalently-bound ligands. Our results show that this approach is able to increase the accuracy of AlphaFold 3 models. We explore the opportunities and limitations of this method, which has been implemented as a proof-of-concept pipeline named AF3x, available at https://github.com/KosinskiLab/af3x.

## Introduction

The field of structural biology has been revolutionized by the advent of deep learning techniques, particularly with the introduction of AlphaFold by DeepMind. AlphaFold2 (Evans et al., 2021; Jumper et al., 2021) set a new benchmark for predicting protein structures with remarkable accuracy, effectively addressing one of the long-standing challenges in biology. Building on this success, AlphaFold 3 (Abramson et al., 2024) has recently expanded the capabilities to model not only proteins but also nucleic acids like DNA and RNA, along with their interactions with small molecules and post-translational modifications.

Despite these advancements, certain limitations remain. AlphaFold 3, like its predecessors, struggles with modeling specific structural conformations or interactions, particularly when evolutionary data is sparse due to a limited number of homologous sequences or weak co-evolutionary signals within a complex. Protein-antibody interactions exemplify this challenge, as they are highly variable, structurally unique, and lack co-evolutionary signals.

Previous studies with AlphaFold2 demonstrated that incorporating experimental data can enhance modeling outcomes of AlphaFold2. Specifically, the use of distance restraints derived from crosslinking mass spectrometry (XL-MS) has shown promise in guiding the folding process toward more accurate conformations (Discovery et al., 2024; Feng et al., 2024; Stahl et al., 2024, 2023; Xie et al., 2024; Zhang et al., 2023). XL-MS is an experimental technique that provides distance constraints between specific amino acid residue types within or between proteins, offering spatial information to complement computational predictions (Graziadei and Rappsilber, 2022).

In XL-MS, crosslinkers—bifunctional chemical reagents—are used to covalently link amino acid residues that are in close proximity. These crosslinkers are designed with reactive groups at both ends, which target specific residue types, typically lysines or other nucleophilic groups, depending on the chemical properties of the crosslinker. The bifunctional nature of the crosslinker ensures that it connects only residues within a certain distance range, typically spanning 0 to 35 Å, depending on spacer length and chemistry of the crosslinker. This creates a permanent covalent bond between the residues. After crosslinking, the protein or protein complex is enzymatically digested, and the resulting peptide fragments are analyzed using MS. Crosslinked peptides are identified based on their unique mass shifts, which correspond to the addition of the crosslinker. Computational tools then map these crosslinks onto protein structures, providing experimentally derived distance restraints that help refine or validate structural models.

Crosslinks are incorporated as restraints in structural modeling by converting the spatial information they provide into distance constraints that guide the placement of residues during model generation. These distance restraints are typically defined as upper bounds, reflecting the maximum distance allowed between the residues linked by the crosslinker. Commonly, harmonic distance restraints are employed, where the potential energy increases quadratically if the modeled distance deviates from the specified range, encouraging the residues to remain within the experimentally determined constraints. Bayesian approaches are also used, where crosslink data are treated as probabilistic evidence, integrated into the modeling process to influence the likelihood of certain conformations (Erzberger et al., 2014; Ferber et al., 2016).

Several computational tools are designed to incorporate such distance restraints into structural modeling workflows. HADDOCK (High Ambiguity Driven biomolecular DOCKing) is widely used for docking and refinement of biomolecular structures using ambiguous and unambiguous distance restraints, including those derived from XL-MS (Cyril Dominguez et al., 2003). IMP (Integrative Modeling Platform) allows for the integration of crosslink data with other experimental information to produce ensemble models of biomolecular complexes (Webb et al., 2018). AlphaLink, GRASP, and other AlphaFold2-based tools (Feng et al., 2024; Stahl et al., 2024, 2023; Xie et al., 2024; Zhang et al., 2023) adapt the AlphaFold2 framework to incorporate crosslink restraints. Chai-1 reproduces AlphaFold 3 network incorporating distance restraints (Discovery et al., 2024).

All these tools represent crosslinks as distance restraints or residue pair representation weighting, an approach that offers significant advantages in terms of computational efficiency and versatility. This approach is compatible with a wide range of optimization algorithms and can be seamlessly integrated into deep learning frameworks, such as by incorporating crosslink information into token pair representations as trainable weight matrices. Additionally, multiple distance restraints can be applied simultaneously to a single residue if it participates in multiple crosslinks, even though, in reality, a single residue can only bind one crosslinker at a time within a molecule. The distance-based approach also facilitates modeling in cases where subsets of crosslinks conflict and cannot be satisfied by a single model, providing a straightforward method to address ambiguities in experimental data. However, this representation has notable limitations. While distance restraints can be parameterized using experimental crosslink length distributions, they fail to account for the steric and conformational constraints imposed by the atomic structure and composition of the crosslinker. This may result in favorable crosslink scores even when the crosslinks are inconsistent with the combined structure of the macromolecule, its surroundings, and the crosslinker.

An alternative to this approach is the explicit representation of crosslinks, which models the full or coarse-grained atomic structure of the crosslinker. Tools like xWalk and Jwalk (Bullock et al., 2016; Kahraman et al., 2011) implement this concept, simulating crosslinks more realistically by considering the shortest path on the protein surface based on structural models. However, the computational demands of such explicit calculations have so far limited their use to scoring rather than serving as direct restraints in modeling workflows (Kahraman et al., 2013).

Explicit representations also have their limitations, such as high computational cost and the restriction that a single residue can only accommodate one crosslinker at a time. Despite these drawbacks, explicit crosslinks provide the benefit of imposing more realistic spatial restraints and accounting for steric exclusion effects from neighboring atoms, potentially improving the physical plausibility of structural models.

In this study, we explore an approach to integrating XL-MS data into AlphaFold 3 by explicitly modeling crosslinks as covalently-bound ligands. This method aims to directly incorporate the experimental distance restraints into the structural prediction process, potentially overcoming some of the limitations associated with modeling complexes lacking extensive evolutionary information.

## Methods

Crosslinkers were incorporated into AlphaFold 3 as custom ligands using the userCCD feature in a custom software code named AF3x. For each crosslinker, the possible bonds to amino acid residues were defined using the bondedAtomPairs parameter. The current implementation supports 13 crosslinkers, including DSSO, DSS, DSG, BS3, and azide-A-DSBSO, BS2G, DSBU, PHOX, BSPEG5, BSPEG9, SDA, LCSDA, and SDAD with additional crosslinkers easily added based on the provided template.

To run the protocol, users need to provide a list of crosslinked residue pairs along the crosslink name and based on that the userCDD and bondedAtomPairs will be added automatically behind the scenes. By default, only one crosslink per residue will be selected by the protocol behind the scenes to ensure consistency with experimental constraints. However, this restriction can be disabled, enabling AlphaFold 3 to determine which crosslink to apply when multiple options exist. False positive crosslinks are not explicitly excluded during modeling. Instead, AlphaFold 3 sometimes automatically resolves conflicts by breaking bonds for crosslinks that cannot be satisfied within the structural model, ensuring that only plausible configurations are retained. Additionally, a procedure is also implemented to sample all possible crosslink combinations of a given subset count, allowing an exploration of potential crosslinking configurations.

All AlphaFold 3 and AF3x runs were run using 20 seeds with five diffusion samples per seed (100 models in total), unless stated otherwise. The sequences databases were downloaded on November 2021 using the script provided in AlphaFold 3. The PDB template database was filtered by setting the maximum template release date to the 1st of January of the year where the true structure was released to exclude the structures from the template set.

For the PDB ID: 9G5K test case, a single simulated azide-A-DSBSO crosslink connecting the residue 104 of SLC19A3 to lysine 43 of the nanobody was used.

For the PDB ID: 4G3Y test case, the following crosslinks obtained using DSSO by (Di Ianni et al., 2024) were used:

| Crosslink ID | Chain1 | AbsPos1 | Chain2 | AbsPos2 | Protein1 | Protein1 |
| --- | --- | --- | --- | --- | --- | --- |
| **1** | **B** | **53** | **C** | **66** | **>sp\|P01857\|Infliximab_HC** | **>sp\|P01375\|TNF_alpha** |
| 2 | B | 54 | C | 66 | >sp\|P01857\|Infliximab_HC | >sp\|P01375\|TNF_alpha |
| **3** | **B** | **54** | **C** | **113** | **>sp\|P01857\|Infliximab_HC** | **>sp\|P01375\|TNF_alpha** |
| 4 | B | 53 | C | 113 | >sp\|P01857\|Infliximab_HC | >sp\|P01375\|TNF_alpha |
| 5 | B | 54 | C | 106 | >sp\|P01857\|Infliximab_HC | >sp\|P01375\|TNF_alpha |
| **6** | **B** | **67** | **C** | **129** | **>sp\|P01857\|Infliximab_HC** | **>sp\|P01375\|TNF_alpha** |
| 7 | B | 208 | C | 129 | >sp\|P01857\|Infliximab_HC | >sp\|P01375\|TNF_alpha |
| **8** | **A** | **1** | **C** | **91** | **>sp\|P0DOX8\|Infliximab_LC** | **>sp\|P01375\|TNF_alpha** |
| 9 | A | 1 | C | 99 | >sp\|P0DOX8\|Infliximab_LC | >sp\|P01375\|TNF_alpha |
| 10 | A | 1 | C | 90 | >sp\|P0DOX8\|Infliximab_LC | >sp\|P01375\|TNF_alpha |
| 11 | A | 49 | C | 113 | >sp\|P0DOX8\|Infliximab_LC | >sp\|P01375\|TNF_alpha |
| 12 | A | 145 | C | 129 | >sp\|P0DOX8\|Infliximab_LC | >sp\|P01375\|TNF_alpha |

The crosslinks retained for modeling, following the automated removal of crosslinks linked to the same residue, are highlighted in bold.

Images were generated using UCSF ChimeraX (Goddard et al., 2018).

## Results

### Implementation

Explicit crosslinks were implemented using the custom bonded ligands feature of AlphaFold 3 (see Methods). This approach allows the modeling of crosslinks as covalently-bound ligands, enabling the incorporation of experimentally derived spatial restraints directly into the structure prediction process.

### Correct complex prediction using a simulated crosslink

To verify whether the explicit crosslink approach is feasible, we selected the structure of the human SLC19A3 transporter in its inward-open state bound to a nanobody (Gabriel et al., 2024) (**Figure 1a**, PDB ID: 9G5K, released on 2024-10-02, after the AlphaFold training cutoff date of 30 September 2021) and an azide-A-DSBSO crosslink simulated based on the structure. Using default AlphaFold 3, the interaction was modeled incorrectly, with the nanobody misaligned and placed away from the interaction site in an incorrect conformation (**Figure 1b**, RMSD: 6.47 Å). However, incorporating a single explicit crosslink resulted in a near-perfect prediction (**Figure 1b**, RMSD: 0.81 Å). Notably, even the side-chain rotamers at the interface were predicted accurately, demonstrating a high level of precision in capturing the interaction details even with just one crosslink (**Figure 1d**). Furthermore, the crosslink-based model achieved higher predicted quality metrics compared to the default prediction (iPTM 0.82 vs 0.26 and higher local pLDDT scores **Figure 1e**). This result indicates that the explicit crosslink procedure is effective in at least one case.

**Figure 1.**
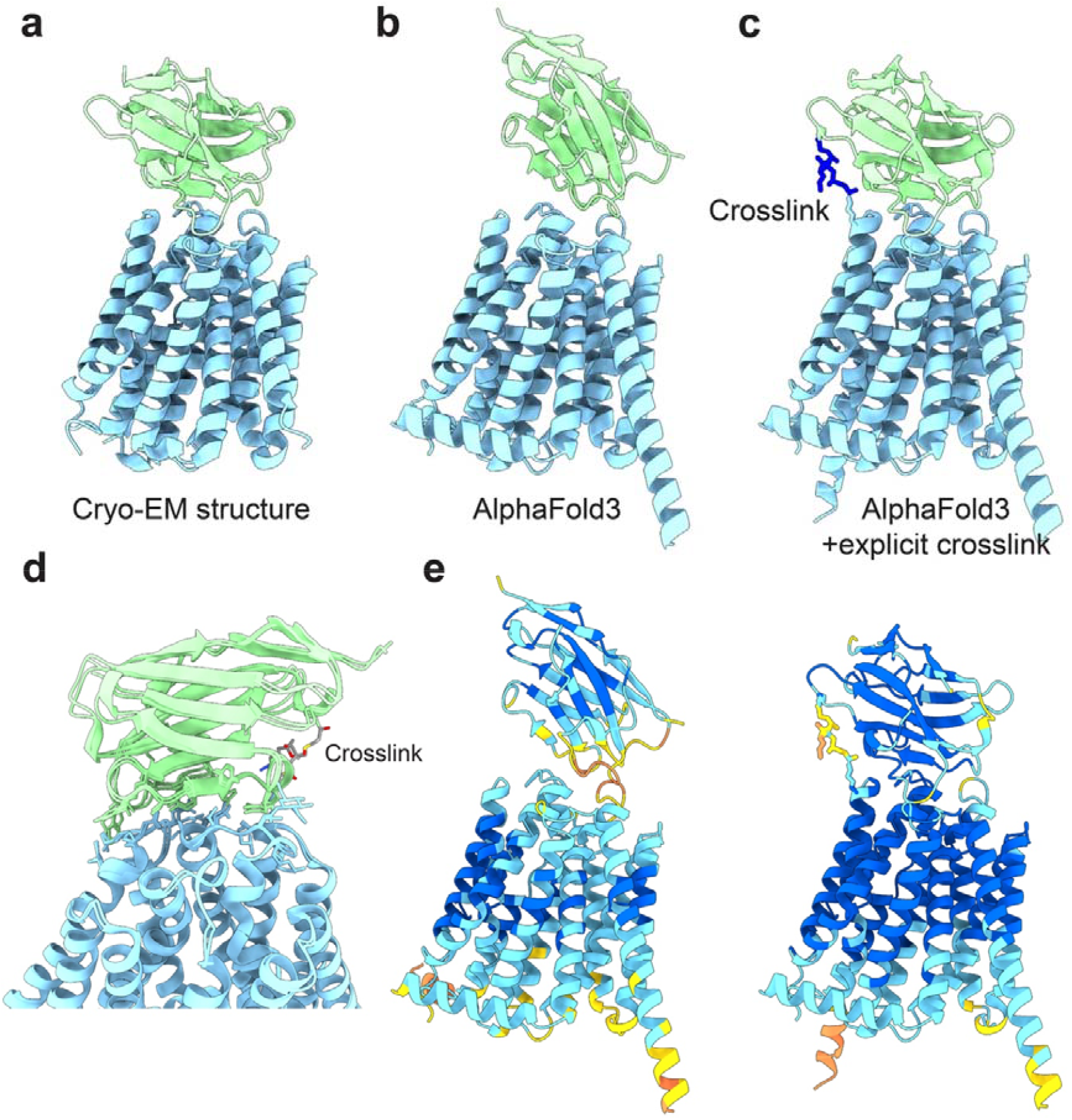
Test on the human SLC19A3 structure in complex with a nanobody. **a.** The cryo-EM structure of the human SLC19A3 transporter in its inward-open state bound to a nanobody (PDB ID: 9G5K), resolved at 3.09 Å resolution. SLC19A3 is colored blue and nanobody is colored salmon. **b.** The AlphaFold 3 model shows the nanobody placed incorrectly, shifted away from the interaction site and in an inaccurate conformation. **c.** The AlphaFold 3 model, generated using the azide-A-DSBSO crosslinker, accurately predicts the interaction, aligning the nanobody correctly at the binding site. **d.** A superposition of the cryo-EM structure and the crosslinked model highlights the close agreement, with interface side chains displayed to demonstrate the accurate prediction of rotamers and interaction details. **e,** The models from b and c colored by each residue’s predicted local distance difference test (pLDDT) score. A residue with a pLDDT score that is greater than 90 (dark blue) indicates high estimated accuracy of the position of its backbone and sidechain rotamers. A pLDDT score above 70 (light blue) suggests that the prediction of the backbone is confident. The disordered regions in the models are removed from display for clarity.

### AlphaFold 3 can deal with conflicting crosslinks

XL-MS data often include conflicting crosslinks, which may arise from false positive identifications, alternative conformations, or protein aggregation. To test whether AlphaFold 3 can handle such conflicts, we selected the crystal structure of TNF-alpha in complex with the Infliximab Fab fragment (Liang et al., 2013) (PDB ID: 4G3Y, released on 2013-03-27), for which experimental crosslinking data is available (Di Ianni et al., 2024). This dataset contains some crosslinks that are incompatible with the crystallized interaction site (**Figure 2**a). As the structure predates the AlphaFold 3 training cutoff, it was likely included in the training dataset. Consequently, AlphaFold 3 predicts this structure nearly perfectly (RMSD: 0.88 Å).

**Figure 2.**
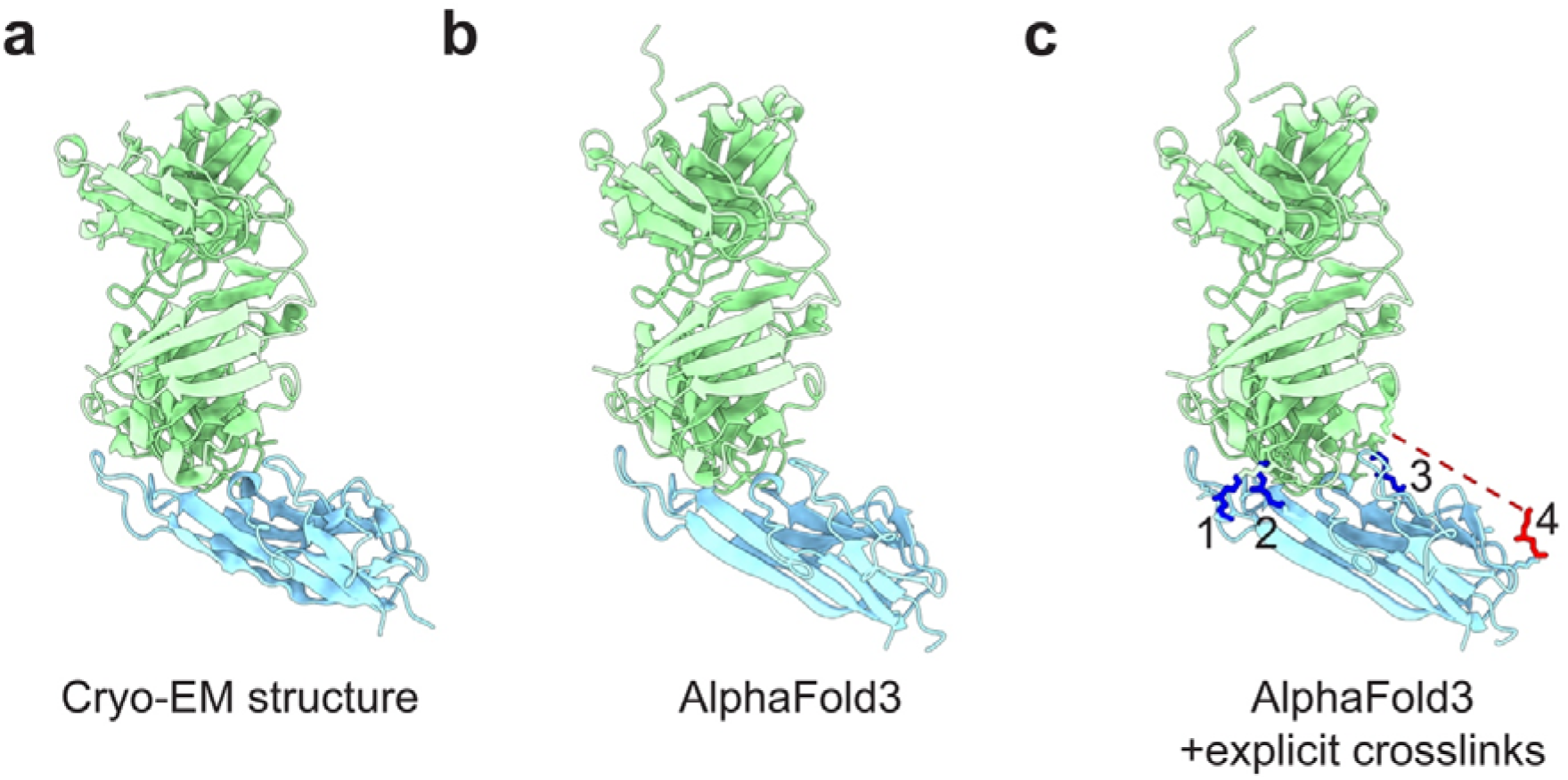
An example of resolving conflicting crosslinks. **a.** The X-ray structure (PDB ID: 4G3Y) resolved at 2.60 Å. **b.** The AlphaFold 3 model. **c.** The AlphaFold 3 model built with crosslinks, with DSSO crosslinks explicitly represented (indicated with numbers). The dashed line represents the broken bond corresponding to the conflicting crosslink. In c and d, disordered regions are removed from display for clarity.

To evaluate the ability of the explicit crosslink protocol to handle conflicting crosslinks, we modeled the structure with explicit crosslinks while enforcing a one-crosslink-per-residue rule. Of the 18 possible crosslinks, four were retained, including one crosslink incompatible with the other three. The resulting model remained close to perfect (RMSD: 1.31 Å), indicating that AlphaFold 3 can successfully manage conflicting crosslinks without being misled, at least in this case.

## Discussion and conclusions

This study demonstrates that incorporating crosslinks as explicit molecules, rather than as distance restraints, improves the AlphaFold 3 modeling accuracy of protein-antibody complexes in certain cases. Although the dataset used was small and limited to protein-antibody complexes, the results suggest that this approach could be a viable XL-MS-based modeling strategy for other systems as well. Testing on larger datasets with real crosslinks and diverse complex types is necessary to evaluate the generalizability of this method. Additionally, the approach could potentially be extended to include crosslinks involving nucleic acids, broadening its applicability. Zero-length crosslinks, such as those created by reagents like EDC and CDI, could also be implemented by replacing standard amino acids with modified versions and defining the linked atoms as covalently bound ligands. While the results indicate that AlphaFold 3 can handle conflicting crosslinks, iterative sampling of different crosslink sets could enhance the robustness of the method. This could be implemented using strategies inspired by semi-Bayesian approaches (Ferber et al., 2016) or the iterative procedures proposed in GRASP (Xie et al., 2024).

Some limitations of this strategy are evident. For example, the method may struggle with residues involved in multiple crosslinks, a situation where distance-based methods, such as HADDOCK, AlphaLink, GRASP, and Chai-1, might be more appropriate. Moreover, the explicit crosslink protocol cannot support interacting residue restraints, which are available in programs like HADDOCK and GRASP. However, excluding some crosslinks and other restraints could serve as a means of validation. Preliminary analyses suggest that even one crosslink can suffice to guide AlphaFold 3 to its global minimum, where its neural network refines the interface, aligning with observations about AlphaFold2 conformational sampling (Roney, 2022).

In conclusion, this work provides a proof of concept that XL-MS-based modeling using explicit crosslinks can improve the accuracy of structural modeling. One could envision a future where other experimental measurements are modeled explicitly, rather than relying on mathematical formulations.

*This manuscript is intended to be updated with additional test examples. Please contact us if you have or wish to contribute a suitable test example. Ideally, the example should 1) fail with standard AlphaFold3 and 2) have real experimental XL-MS data available.*

## Acknowledgements

I thank Agnieszka Obarska-Kosinska for the idea and help in preparing the crosslink definitions. The more polished sections of the code, if any, were written with the assistance of ChatGPT and GitHub Copilot. ChatGPT was also used to draft and refine this manuscript. The selection of the 4G3Y as a test case was informed by the GRASP method manuscript.

## Data and code availability

The code and test data used in this manuscript are available at https://github.com/KosinskiLab/af3x. Note that most of the code and models generated with AF3x are subject to the original AlphaFold 3 license. The crosslink definition file (crosslink_definitions.py) is provided under the GNU General Public License Version 3.

## References

Abramson, J., Adler, J., Dunger, J., Evans, R., Green, T., Pritzel, A., Ronneberger, O., Willmore, L., Ballard, A.J., Bambrick, J., Bodenstein, S.W., Evans, D.A., Hung, C.-C., O’Neill, M., Reiman, D., Tunyasuvunakool, K., Wu, Z., Žemgulytė, A., Arvaniti, E., Beattie, C., Bertolli, O., Bridgland, A., Cherepanov, A., Congreve, M., Cowen-Rivers, A.I., Cowie, A., Figurnov, M., Fuchs, F.B., Gladman, H., Jain, R., Khan, Y.A., Low, C.M.R., Perlin, K., Potapenko, A., Savy, P., Singh, S., Stecula, A., Thillaisundaram, A., Tong, C., Yakneen, S., Zhong, E.D., Zielinski, M., Žídek, A., Bapst, V., Kohli, P., Jaderberg, M., Hassabis, D., Jumper, J.M., 2024. Accurate structure prediction of biomolecular interactions with AlphaFold 3. Nature 1–3. 10.1038/s41586-024-07487-w

Bullock, J.M.A., Schwab, J., Thalassinos, K., Topf, M., 2016. The Importance of Non-accessible Crosslinks and Solvent Accessible Surface Distance in Modeling Proteins with Restraints From Crosslinking Mass Spectrometry *. Molecular & Cellular Proteomics 15, 2491–2500. 10.1074/mcp.M116.058560

Cyril Dominguez, Rolf Boelens, and, Bonvin*, A.M.J.J., 2003. HADDOCK: A Protein−Protein Docking Approach Based on Biochemical or Biophysical Information [WWW Document]. ACS Publications. 10.1021/ja026939x

Di Ianni, Andrea, Di Ianni, Alessio, Cowan, K., Barbero, L.M., Sirtori, F.R., 2024. Leveraging Cross-Linking Mass Spectrometry for Modeling Antibody–Antigen Complexes. J. Proteome Res. 23, 1049–1061. 10.1021/acs.jproteome.3c00816

Discovery, C., Boitreaud, J., Dent, J., McPartlon, M., Meier, J., Reis, V., Rogozhnikov, A., Wu, K., 2024. Chai-1: Decoding the molecular interactions of life. 10.1101/2024.10.10.615955

Erzberger, J.P., Stengel, F., Pellarin, R., Zhang, S., Schaefer, T., Aylett, C.H.S., Cimermančič, P., Boehringer, D., Sali, A., Aebersold, R., Ban, N., 2014. Molecular Architecture of the 40S_eIF1_eIF3 Translation Initiation Complex. Cell 158, 1123–1135. 10.1016/j.cell.2014.07.044

Evans, R., O’neill, M., Pritzel, A., Antropova, N., Senior, A., Green, T., Žídek, A., Bates, R., Blackwell, S., Yim, J., Ronneberger, O., Bodenstein, S., Zielinski, M., Bridgland, A., Potapenko, A., Cowie, A., Tunyasuvunakool, K., Jain, R., Clancy, E., Kohli, P., Jumper, J., Hassabis, D., 2021. Protein complex prediction with AlphaFold-Multimer. bioRxiv 2021.10.04.463034. 10.1101/2021.10.04.463034

Feng, S., Chen, Z., Zhang, C., Xie, Y., Ovchinnikov, S., Gao, Y.Q., Liu, S., 2024. Integrated structure prediction of protein–protein docking with experimental restraints using ColabDock. Nat Mach Intell 6, 924–935. 10.1038/s42256-024-00873-z

Ferber, M., Kosinski, J., Ori, A., Rashid, U.J., Moreno-Morcillo, M., Simon, B., Bouvier, G., Batista, P.R., Müller, C.W., Beck, M., Nilges, M., 2016. Automated structure modeling of large protein assemblies using crosslinks as distance restraints. Nature methods 13, 515–20. 10.1038/nmeth.3838

Gabriel, F., Spriestersbach, L., Fuhrmann, A., Jungnickel, K.E.J., Mostafavi, S., Pardon, E., Steyaert, J., Löw, C., 2024. Structural basis of thiamine transport and drug recognition by SLC19A3. Nat Commun 15, 1–13. 10.1038/s41467-024-52872-8

Goddard, T.D., Huang, C.C., Meng, E.C., Pettersen, E.F., Couch, G.S., Morris, J.H., Ferrin, T.E., 2018. UCSF ChimeraX: Meeting modern challenges in visualization and analysis. Protein Science 27, 14–25. 10.1002/pro.3235

Graziadei, A., Rappsilber, J., 2022. Leveraging crosslinking mass spectrometry in structural and cell biology. Structure 30, 37–54. 10.1016/j.str.2021.11.007

Jumper, J., Evans, R., Pritzel, A., Green, T., Figurnov, M., Ronneberger, O., Tunyasuvunakool, K., Bates, R., Žídek, A., Potapenko, A., Bridgland, A., Meyer, C., Kohl, S.A.A., Ballard, A.J., Cowie, A., Romera-Paredes, B., Nikolov, S., Jain, R., Adler, J., Back, T., Petersen, S., Reiman, D., Clancy, E., Zielinski, M., Steinegger, M., Pacholska, M., Berghammer, T., Bodenstein, S., Silver, D., Vinyals, O., Senior, A.W., Kavukcuoglu, K., Kohli, P., Hassabis, D., 2021. Highly accurate protein structure prediction with AlphaFold. Nature 596, 583– 589. 10.1038/s41586-021-03819-2

Kahraman, A., Herzog, F., Leitner, A., Rosenberger, G., Aebersold, R., Malmström, L., 2013. Cross-Link Guided Molecular Modeling with ROSETTA. PLOS ONE 8, e73411. 10.1371/journal.pone.0073411

Kahraman, A., Malmström, L., Aebersold, R., 2011. Xwalk: computing and visualizing distances in cross-linking experiments. Bioinformatics (Oxford, England) 27, 2163–4. 10.1093/bioinformatics/btr348

Liang, S., Dai, J., Hou, S., Su, L., Zhang, D., Guo, H., Hu, S., Wang, H., Rao, Z., Guo, Y., Lou, Z., 2013. Structural Basis for Treating Tumor Necrosis Factor α (TNFα)-associated Diseases with the Therapeutic Antibody Infliximab*. Journal of Biological Chemistry 288, 13799–13807. 10.1074/jbc.M112.433961

Roney, J.P., 2022. State-of-the-Art Estimation of Protein Model Accuracy Using AlphaFold. Phys. Rev. Lett. 129. 10.1103/PhysRevLett.129.238101

Stahl, K., Graziadei, A., Dau, T., Brock, O., Rappsilber, J., 2023. Protein structure prediction with in-cell photo-crosslinking mass spectrometry and deep learning. Nat Biotechnol 41, 1810–1819. 10.1038/s41587-023-01704-z

Stahl, K., Warneke, R., Demann, L., Bremenkamp, R., Hormes, B., Brock, O., Stülke, J., Rappsilber, J., 2024. Modelling protein complexes with crosslinking mass spectrometry and deep learning. Nat Commun 15, 7866. 10.1038/s41467-024-51771-2

Webb, B., Viswanath, S., Bonomi, M., Pellarin, R., Greenberg, C.H., Saltzberg, D., Sali, A., 2018. Integrative structure modeling with the Integrative Modeling Platform. Protein Science 27, 245–258. 10.1002/pro.3311

Xie, Y., Zhang, C., Li, S., Du, X., Wang, M., Hu, Y., Liu, S., Gao, Y.Q., 2024. Integrating various Experimental Information to Assist Protein Complex Structure Prediction by GRASP. 10.1101/2024.09.16.613256

Zhang, Y., Zhang, Z., Kagaya, Y., Terashi, G., Zhao, B., Xiong, Y., Kihara, D., 2023. Distance-AF: Modifying Predicted Protein Structure Models by Alphafold2 with User-Specified Distance Constraints. 10.1101/2023.12.01.569498

